# Looking beyond stereotyped neuron structures reveals links between beading and morphological rearrangements in aging phenotypes

**DOI:** 10.64898/2026.06.15.732273

**Authors:** Kin Gomez, Kyle Nguyen, John Lagergren, Kevin Flores, Adriana San Miguel

## Abstract

Understanding how neuronal morphology changes during aging and acute stress is essential for elucidating mechanisms of neurodegeneration. The highly branched PVD neuron of *Caenorhabditis elegans* provides a powerful model for studying dendritic remodeling and degeneration-associated phenotypes such as dendritic beading. However, the complexity of this arbor presents substantial challenges for automated segmentation and quantitative analysis. In this study, we adapted a convolutional neural network (CNN)-guided region growing framework for automated dendrite tracing, coupled with two topology-based algorithms for categorizing dendritic segments by branching degree. The segmentation algorithm achieved high accuracy relative to manual tracing, with a median Dice coefficient of 0.82, while reducing analysis time by approximately tenfold. Automated dendrite categorization demonstrated strong agreement with manual annotations across branching orders, though position-based mapping performance declined with age due to progressive morphological distortion. Leveraging this platform, we investigated mechanistic differences in dendritic beading patterns observed during aging and cold shock. Consistent with prior work, aging was associated with decreased inter-bead spacing, whereas cold shock produced increased bead dispersion with stress severity. Structural analysis revealed that these trends were not driven by dendritic pruning or reduced arbor complexity. Instead, while a traditional anatomically unflexible paradigm falsely implicated lower-degree dendrites as highly vulnerable, our branching-informed framework revealed that age-dependent beading is fundamentally dictated by a segment’s history of successive branching events. Conversely, acute cold shock triggered systemic beading that expanded across all dendritic orders in a severity-dependent manner. Together, these findings demonstrate that chronic aging and acute stress engage distinct degenerative pathways (compartment-specific lineage vulnerability versus global architectural collapse) rather than gross morphological loss, as well as highlighting the need for paradigms that enable reliable analysis of changing morphologies.

## 2 Introduction

Understanding neuronal morphology is critical for uncovering how neurodegenerative conditions and acute pathological episodes alter cellular phenotypes. Progressive alterations in branching complexity, dendritic length, and spine density have been widely documented across standard models of Alzheimer’s Disease, Parkinson’s Disease, and Amyotrophic Lateral Sclerosis, establishing a direct link between structural disruption and impaired synaptic connectivity [1–4]. Quantifying these subtle structural changes requires highly precise segmentation of neuronal processes. However, neurons often exhibit elaborate, multi-tiered architectures that create significant bottlenecks for automated analysis platforms. The *Caenorhabditis elegans* PVD somatosensory neuron offers an exceptional model for resolving these architectural dynamics due to its highly stereotyped, "menorah-like" dendritic arbor [5]. This intricate branching hierarchy is linked to function, structural disorganization of the menorah units directly reduces mechanosensory capability [6], and the arbor is susceptible to degenerative beading phenotypes driven by aging and acute environmental stress [7,8]. Yet, while macro-level morphological shifts have been observed under these stressors, a high-resolution, quantitative description of degenerative patterns relative to specific branch positions within the menorah structure has remained completely unknown.

Detailed analysis of neuronal arbors enables the extraction of essential morphological metrics, such as Sholl-based branching complexity, and provides the necessary spatial framework to track labeled subcellular structures across cellular compartments. Existing computational tools, such as the Fiji Simple Neurite Tracer (SNT) add-on, are widely utilized for manual or semi-automated tracing [9]. While these conventional approaches provide operational flexibility, they remain highly lab or-intensive and susceptible to operator bias. Furthermore, the extreme structural complexity of thin, dense dendritic arbors (while biologically relevant) creates substantial segmentation bottlenecks for traditional intensity- or threshold-based tracking algorithms. This challenge is severely compounded in crowded imaging fields or when analyzing multi-channel datasets, prompting previous engineering efforts to look toward Convolutional Neural Networks (CNNs) to achieve robust segmentation [10,11]. A pioneering milestone in this domain was recently achieved by Yuval et al., who developed an elegant pipeline combining CNN-based classification with algorithmic categorization of the canonical PVD menorah structure [12]. This framework demonstrated the power of deep learning to uncover subtle, high-precision phenotypical variations and structural motifs in the PVD arbor, such as symmetrical three-way junctions. However, because their categorization framework is optimized specifically for well-organized, intact PVD architectures, it struggles to adapt efficiently to the structural degeneration inherent to aging and stress. This limitation leaves a critical technical gap, highlighting the need for an accelerated, structure-agnostic pipeline capable of handling the high sample volumes required for population-level screening.

To address these complex, pixel-level tracking limitations, computer vision frameworks have increasingly shifted away from standalone semantic segmentation architectures toward hybrid pipelines. Historically, artificial neural networks were confined to post-processing roles or used primarily to predict static thresholds for traditional region growing techniques [13,14]. However, recent hybrid paradigms train convolutional neural networks (CNNs) to output dense probability maps that directly govern pixel inclusion, allowing localized classification to enforce global spatial context. When compared against conventional architectures like U-Nets, these hybrid CNN-guided region growing algorithms have demonstrated superior performance in preserving thin, continuous, and topologically intricate features, achieving exceptional pixel-level precision in structural analogues like leaf veins and retinal vascular networks [10,11].

In this study, we present a neurite tracing and classification tool specifically optimized for the PVD neuron structure. Our method leverages a specialized CNN-based region growing algorithm [10,11], coupled with topology-based branch classification to achieve improved segmentation accuracy relative to expert manual annotation, while drastically reducing user intervention time compared to manual tools like Fiji’s SNT. Furthermore, the tool offers two distinct modes for categorizing dendritic processes: a canonical framework based on segment positions within the stereotyped menorah architecture, and a structure-agnostic framework that classifies segments sequentially based strictly on local branching events. We demonstrate that this second, branching-centric approach uncovers critical morphometric insights regarding the progression of age-related dendritic beading that are otherwise obscured when using a traditional, menorah-aware paradigm.

## 3 Materials and Methods

### 3.1 Worm culture and imaging

We conducted experiments using the *C. elegans* strain NC1686 (wdIs51 [F49H12.4::GFP + unc-119(+)]), a wild type background expressing GFP in the PVD neuron. Animals were synchronized by bleaching (solution of 1 M NaOH, sodium hypochlorite, and deionized water at a 2:1:1 ratio) and maintained at 20 °C on nematode growth medium (NGM) agar plates seeded with E. coli OP50 until adulthood. Adult worms were transferred with a platinum wire onto 2% agarose pads prepared on glass slides. For immobilization, a drop of 2 mM tetramisole in M9 buffer was applied before placing a coverslip on top. For the aging dataset, worms were transferred to fresh plates on non-imaging days to maintain adequate feeding.

Imaging was performed with a Leica DMi8 equipped with a CrestOptics X-light V2 spinning disk confocal module and a Hamamatsu Orca-Fusion camera, using a 63× objective. Samples were illuminated with an 89North laser diode illumination system. Acquisition conditions were kept constant across experiments (60ms exposure, 30% laser power). Z-stacks of 30 images at 1 μm spacing were collected, and maximum-intensity projections were generated for downstream analysis.

### 3.2 Bead detection

The determination of bead instances along the dendrites was performed using the automated bead detection pipeline introduced by Sahand et al. (2020) [8]. Prior to bead identification, TIFF images were converted to 8-bit PNG format using a percentile-based intensity normalization procedure. For each image, pixel intensities were rescaled using the 1st and 99th percentile values as the lower and upper intensity bounds, respectively. Intensities at or below the 1st percentile were mapped to 0, and intensities at or above the 99th percentile were mapped to 255. Intermediate values were linearly scaled between these bounds. An optional brightness multiplier was then applied to the normalized image, with values clipped to the valid 8-bit range before saving. Only objects with a probability of at least 0.7 were kept, and the resulting masks were manually screened for errors before including them in the dataset.

### 3.3 Region Growing effectively distinguishes PVD neurite from background pixels

The CNN-based region growing framework was implemented as a two-step process: (i) training a convolutional neural network (CNN) to predict the probability of foreground (dendrite) versus background pixels, and (ii) recursively applying a region growing algorithm to expand the predicted dendrite segmentation [10,11]. We used a dataset of 27 grayscale images of *C. elegans* dendrites with labels generated using a custom MATLAB-based annotation software. The raw images had dimensions of 2304 × 2304 pixels. Each image was padded with 24 pixels on all sides, resulting in an effective size of 2352×2352 pixels. Pixel intensities were normalized to the range [0,1]. The dataset was split into 21 training images and 6 validation images to enable supervised learning with an independent evaluation set. Each image was divided into tiles of 48×48 pixels. Image tiles were generated by centering on ground-truth dendrite pixels (foreground), and an additional set of background centered tiles was sampled at a 3:1 background-to-foreground ratio. These tiles formed the inputs to the CNN (Figure 1). For each 48 × 48 input tile, the CNN produces a 3 × 3 × 2 classification map covering the center pixel and its eight neighbors. The first channel represents foreground probabilities, while the second channel represents background probabilities for the center pixel and its adjacent neighbor pixels. This dual-channel approach enables simultaneous classification of a local neighborhood rather than just a single pixel, improving robustness in low-contrast regions and later facilitating the region growing algorithm. The network consists of five convolutional blocks that progressively down sample features from the input tile to the 3×3×2 output. Each block contains three convolutional layers, except for the first block, which consists of two convolutional layers. Each convolutional layer is followed by batch normalization, dropout, and a LeakyReLU activation. A Softmax activation is applied at the output to result in properly normalized probabilities. We trained the CNN using a pixel-wise cross-entropy (PCE) loss. Optimization was performed with the Adam algorithm [15], using a learning rate of 1e−3, a batch size of 128, and 10 training epochs.

**Figure 1.**
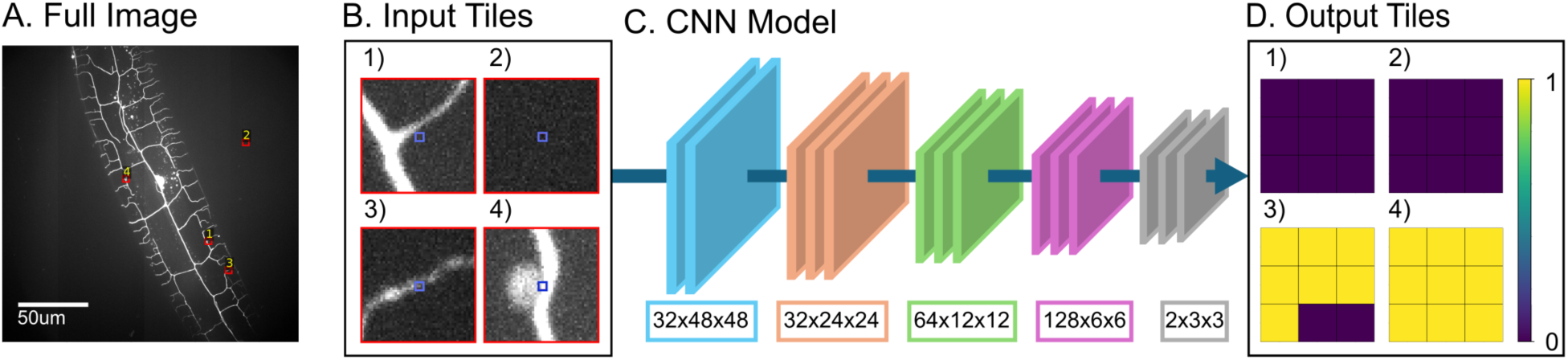
Overview of the CNN-based region growing pipeline. (A) Full grayscale image of *C. elegans* dendrites, with red boxes indicating locations of sampled tiles (red squares). (B) Example 48 × 48 input tiles centered on seed pixels (blue squares). (C) Convolutional neural network architecture: input tiles are progressively down sampled through convolutional blocks (feature map sizes indicated below each block) to generate compact feature representations. (D) Output tiles: for each input tile, the CNN predicts a 3 × 3 × 2 classification map, where each element represents the probability of foreground (dendrite) versus background for the center pixel and its eight neighbors. The output probabilities enforce spatial context and guide the iterative region growing algorithm for dendrite segmentation.

After training, segmentation proceeds via an iterative region growing algorithm as shown in Figure 2. The procedure is initialized by randomly sampling 105 seed pixels within the image. For each seed, a 48 × 48 tile is passed through the CNN to obtain probability estimates for the center pixel and its eight neighbors. Neighboring pixels for which the foreground probability exceeds the background probability by a predefined threshold are appended to the seed list for the next iteration. In this case, we set the threshold to −0.2. It is important to note that this threshold is only used to determine whether a pixel should be added to the considering list, and not to indicate whether the pixel is a foreground pixel (i.e., a dendrite pixel). Once the seed pixels have been considered, they are removed from the considering list. This recursive process continues until convergence (i.e., no new pixels are added), thereby enforcing spatial contiguity and preserving biologically realistic dendritic morphology. Pixels with a foreground probability greater than the background probability are assigned to the segmentation mask. At the end of the iterative region growing process, all pixels classified as foreground (i.e., dendrite pixels) are aggregated to form a comprehensive dendrite mask. This entire two-step process was performed on a single NVIDIA GeForce RTX 4070 GPU.

**Figure 2.**
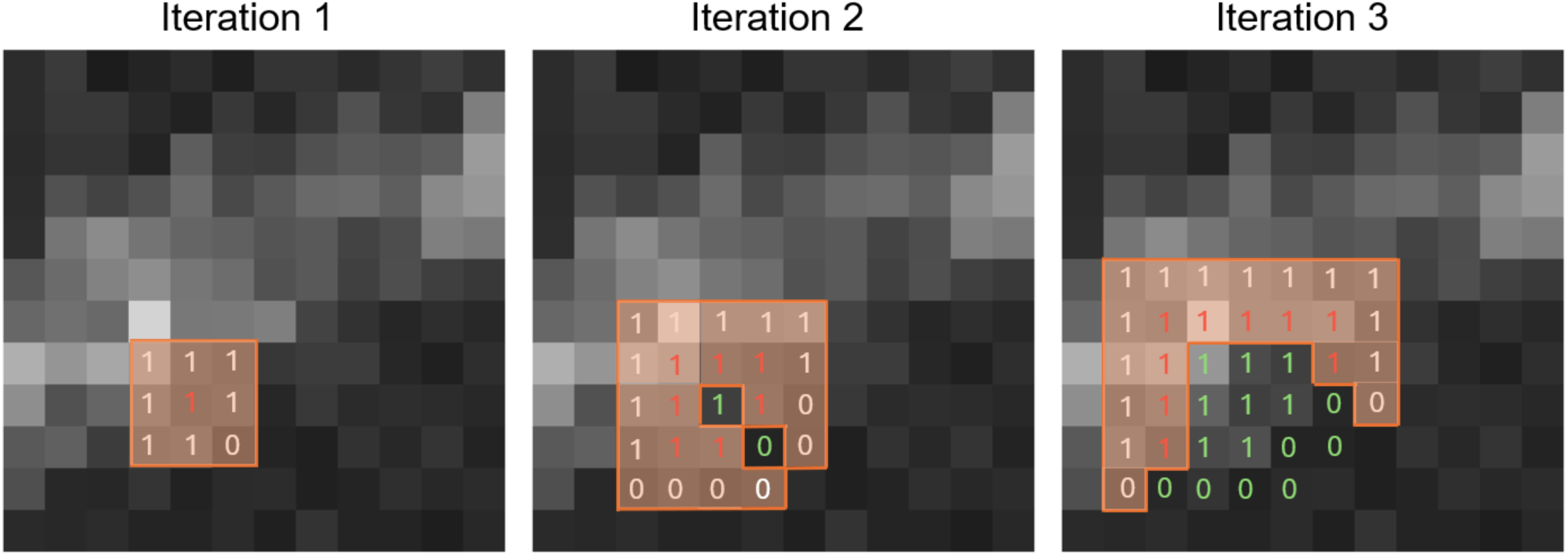
Illustration of the CNN-based region growing process across successive iterations. At each iteration, a seed pixel is selected and its tile is passed through the CNN to predict a 3 × 3 × 2 neighborhood of class probabilities. Foreground predictions (denoted by 1) are appended as new seeds, while background predictions (denoted by 0) are excluded. Iteration 1 begins with a single seed and produces initial foreground classifications for the center pixel (red) and its neighboring pixels (white). The orange box indicates the set of pixels under consideration during each iteration. In Iteration 2, the region expands by adding newly classified foreground neighbors to the seed list, while pixels that have already been considered (green) are removed. Iteration 3 demonstrates further growth, as the region continues to spread along the dendrite while preserving spatial contiguity. The process repeats until no additional foreground pixels are added, resulting in a stable segmentation mask.

### 3.4 Dendrite segment categorization pipeline

After segmentation, we first apply an automated pipeline to categorize dendritic segments by branching degree based on the previously developed algorithm by Yuval et. Al [12] that performs categorization based on the stereotypical PVD menorah structure (Figure 3). Starting from the region growing mask, we perform a preprocessing step to remove false-positive branches that do not belong to the main arbor (Figure 3A). The cleaned mask is skeletonized, and branching points and endpoints are identified. These landmarks delineate the neuronal arbor and allow us to recover the overall orientation and curvature of the worm without requiring a bright-field image of the cuticle. To establish a local coordinate system, splines are fit to the upper and lower boundaries of the arbor, and the centerline is defined as the average of these boundaries (Figure 3B). The tangent of the centerline at each segment centroid serves as the local orientation reference. The skeleton is then partitioned into discrete segments by introducing cuts at each branchpoint. Each cut is drawn along the bisector of the angle between the two daughter branches, ensuring minimal and consistent separation (Figure 3C). Segments are classified as axial if they align within 45° of the local tangent of the centerline, or as radial if they extend outward (Figure 3D).

**Figure 3.**
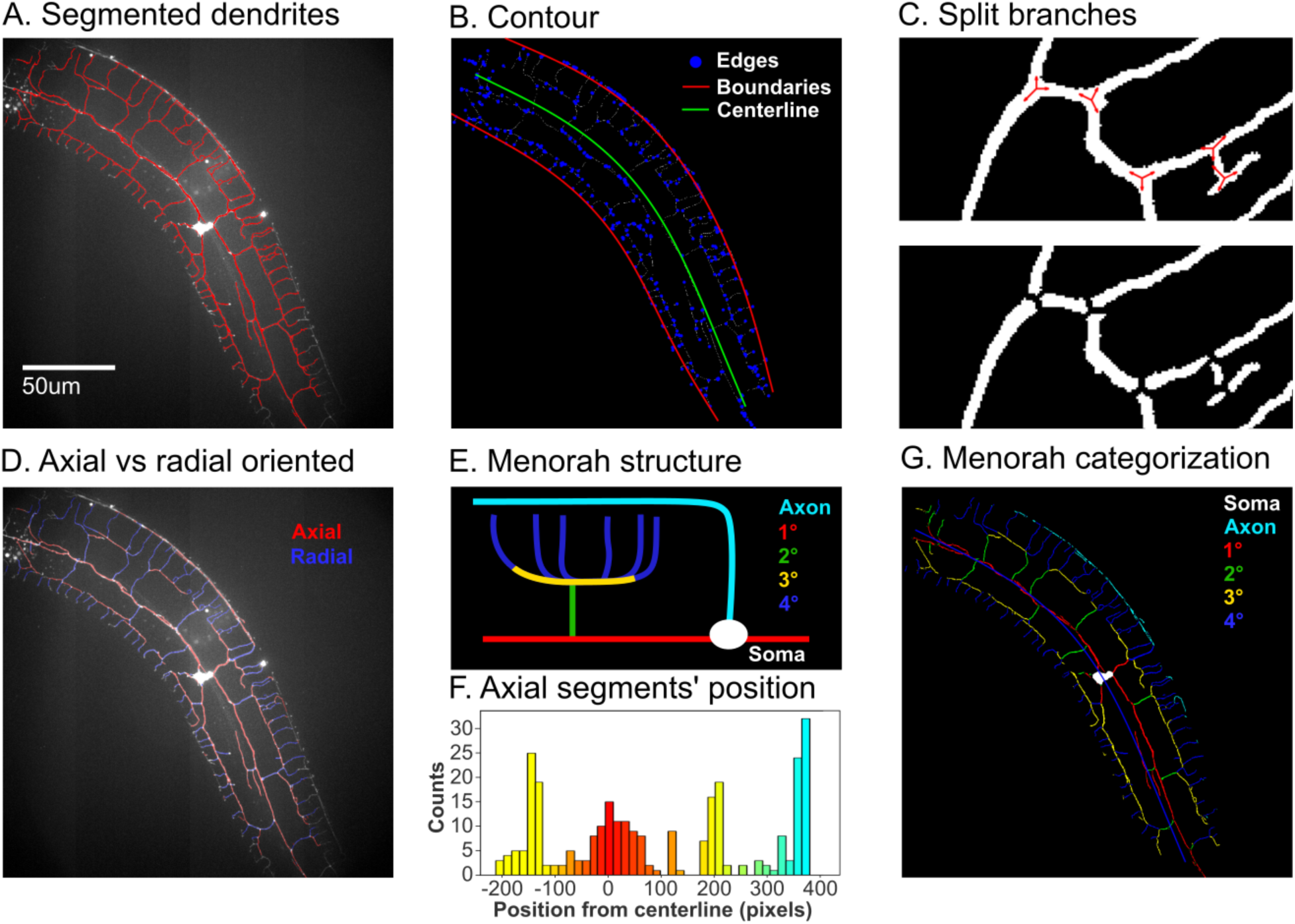
Workflow for categorization of PVD dendrite segments by menorah structure. (A) Example of a segmented section of a PVD neuron. (B) Upper and lower boundaries are extracted by fitting curves along the skeletonized mask, with the centerline defined as their midpoint. (C) Dendrites are split into segments by erasing lines at the bisecting angle of each branch-point. (D) Segments are classified as axial or radial according to their orientation relative to the local centerline tangent. (E) Canonical menorah structure of the PVD neuron. (F) Axial segments are grouped by distance from the centerline into first-degree, third-degree, and axon categories. (G) Radial segments are categorized based on their relative position to adjacent first- and third-degree branches.

Axial segments are further categorized by their distance from the centerline. Owing to the characteristic “menorah” morphology of the PVD arbor, axial elements cluster into distinct bands (Figure 3E-F). We implement a semi-constrained k-nearest-neighbor (KNN) procedure: the first-degree dendrites are anchored at the centerline, and a 3-group KNN is first applied to all axial segments based on signed distance. The two non-centerline clusters are then evaluated for bimodality. If both are unimodal, the 3-group solution is retained. If one cluster is bimodal, the algorithm reruns a 4-group KNN with the centerline class fixed, splitting the bimodal cluster into axon and third-degree categories. In all cases, segments nearest the centerline are assigned as first-degree dendrites, the most medial band (when present) as the axon, and the remaining bands as third-degree dendrites. Radial segments are categorized according to their position relative to neighboring axial branches (Figure 3G). To account for local variability in third-degree distances, the image is subdivided into sections. Radial elements located below the local third-degree average are assigned as second-degree dendrites, whereas those above are assigned as fourth-degree dendrites.

This framework enables robust, reproducible categorization of dendritic segments, even in cases where connectivity is incomplete or fluorescence expression is uneven. By standardizing degree assignment independently of global continuity, the approach facilitates consistent quantification of branching patterns and orientation-dependent phenotypes across different neurons. However, a limitation of this method is that all information regarding parent-child relationships is lost, and any deviation from the stereotypical menorah structure creates artificial branching degree assignment. Consequently, the algorithm is prevented from categorizing any structure beyond the fourth degree of branching, a characteristic that is commonplace in aged neurites or worms harboring loss-of-function mutations such as *eff-1* and *sax-1* [6,16]. To capture this missing information, we developed a secondary dendrite categorization strategy that remains agnostic to the menorah structure and is therefore well-suited for non-canonical PVD organization. Regardless of morphological changes caused by branching, spines, or irregular growth, the primary branch of the PVD neuron consistently serves as the source of all subsequent branches. Even if this principal branch undergoes disruption (such as a change in shape or a pruning event) it retains its status as the foundational branch that all others arise from. Therefore, the first step in our workflow was to identify this structure within the dendrite. To achieve this, we applied Dijkstra’s algorithm to the skeletonized image to find the longest continuous path, which almost always corresponds to the primary (first-degree) branch, unless a high degree of overlap exists between neighboring menorah structures. To mitigate this issue, the soma can be explicitly specified to anchor the path to the main branch. This path is further refined by identifying and eliminating instances of pronounced turns (Figure 4A), the main path identified by Dijkstra’s algorithm is highlighted in yellow, while the blue and red regions are discarded due to sharp turning.

**Figure 4.**
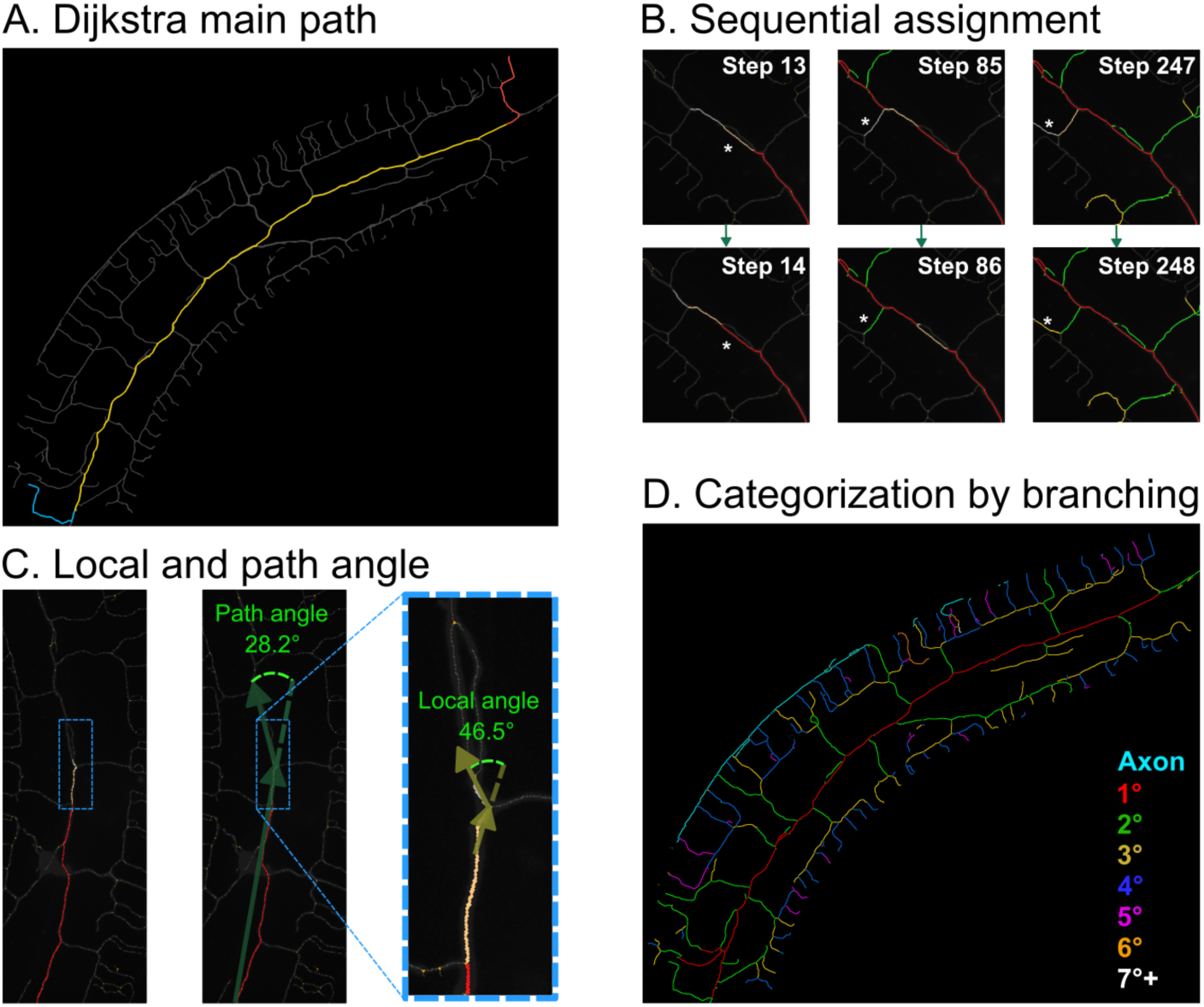
Workflow for categorization of PVD dendrite segments by sequential branching. (A) The main dendrite branch is identified using Dijkstra (yellow), and the twisting edges (red & cyan) are discarded. (B) Example of decision-making events [*] for first, second- and third-degree branches (parent in orange and child in grey). (C) Example of local and path angle between parent and child at different sides of the threshold. (D) Final branching degree assignment.

Once the main path is established, the algorithm sequentially categorizes the regions (divided by junctions), starting from the first element of the main path and moving toward the child segment most likely to belong to the same branch (Figure 4B). All elements that coincide with the previously identified main path are assigned continuity along the first-degree branch. Branching events are evaluated based on three factors: the local angle, the path angle, and the continuity score. The local angle measures the angle of incidence at the junction node between the parent and its children to determine local segmental continuity. Because the organic waviness of neurites often renders local angles insufficient on their own, the path angle calculates the trajectory of all previous elements assigned to a given category against a look-ahead window of the path being evaluated. These two angles (Figure 4C) are averaged to derive a decision angle, which is then compared against a predefined threshold (40°) to determine whether the path should continue the current branch or be categorized as a child of the incoming branch.

Every segment is evaluated by exploring the children of each node while prioritizing lower-degree assignments; specifically, second-degree branches are assigned only after all first-degree branches have been fully determined (Figure 4B). Subsequently, the children of second-degree branches are explored before third-degree assignments begin, and this sequential process continues for higher orders). This hierarchical routing prevents loops from being prioritized and avoids incorrectly assigning high branch numbers to segments in direct contact with low-level branches. Ultimately, this approach yields a more accurate representation of the true parent-child relationships between dendritic segments not being limited to a maximum branching degree (Figure 4D).

## 4 Results

### 4.1 Automated Dendrite Tracing of *C. elegans* with CNN-based Region Growing

We evaluate the performance of the CNN-based region growing algorithm by collecting a total of 46 new images that were not used for training or validation. In Figure 5, we present an example result of a traced mask using the CNN-based region growing algorithm. The left image shows the original image (i.e., input image) of a *C. elegans*. The middle image provides a comparison between the ground truth manually traced mask, created using our MATLAB annotation software, and the CNN-based region growing traced mask. In this comparison, red pixels indicate agreement between the ground truth and our algorithm, green pixels represent traced pixels identified by the algorithm that were not manually traced (i.e., false positives), and blue pixels are those that were manually traced but missed by the algorithm (i.e., false negatives). The right image displays the CNN-based traced mask overlayed on the *C. elegans* image.

**Figure 5.**
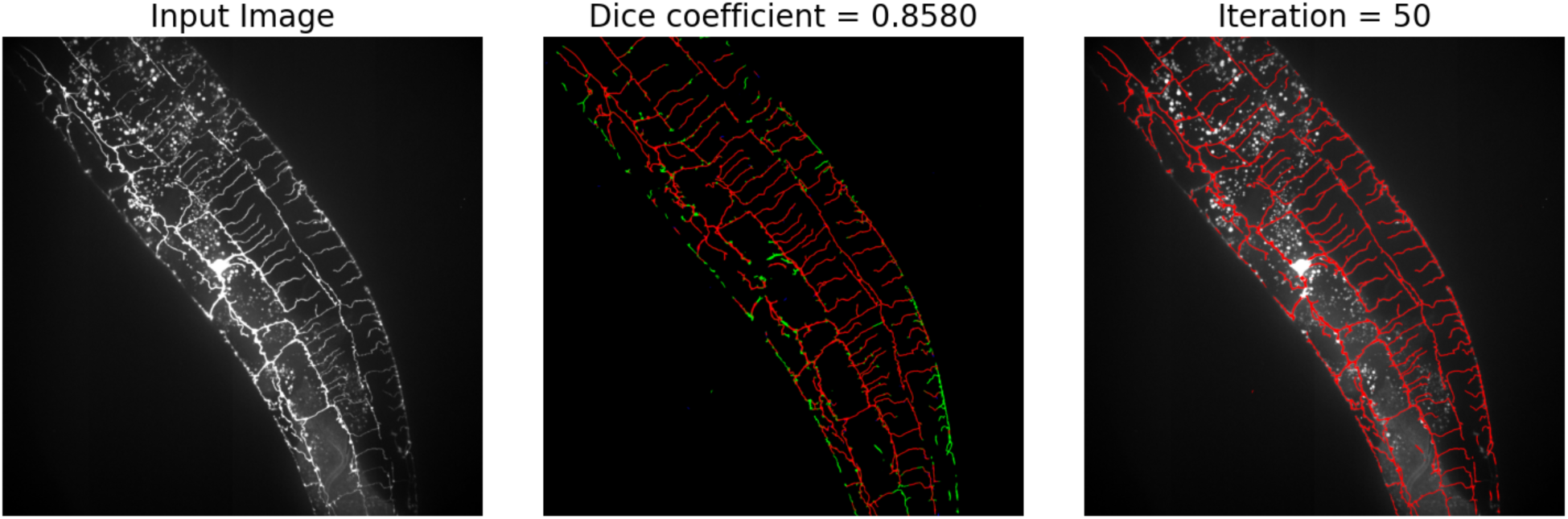
Example result of CNN-based region growing segmentation on a *C. elegans* dendrite image. The left panel shows the original input image. The middle panel compares the manually traced ground-truth mask with the CNN-based region growing output. In this comparison, red pixels indicate agreement between the algorithm and the ground truth, green pixels denote false positives, and blue pixels mark false negatives. The Dice coefficient for this example is 0.858. The right panel overlays the CNN-based segmentation mask (red) on the input image after 50 iterations, illustrating how the algorithm traces dendritic structures.

To assess the performance of the CNN-based region growing algorithm, we compute the Dice score (or Dice similarity coefficient) [17], which measures the overlap between two datasets, typically binary masks in segmentation tasks, ranging from 0 (no overlap) to 1 (perfect overlap). In this example, the Dice score is 0.858 (Figure 5). In Table 1, we display the computed first, second (i.e., median), and third quintiles for the Dice scores as well as the Jaccard Index[18], another related metric for segmentation tasks [19], across different *C. elegans* images in the testing set. A Dice score above 0.7 is often considered acceptable or good in segmentation tasks depending on application area [20–23]. The median Dice score is 0.8235, with the first quintile at 0.7647, indicating a high accuracy in pixel tracing by our algorithm.

**Table 1.** Dice and Jaccard Scores with Q1, Median, and Q3.

| Metric | Q1 | Median | Q3 |
| --- | --- | --- | --- |
| Dice Score | 0.7647 | 0.8235 | 0.8589 |
| Jaccard Index | 0.6191 | 0.6999 | 0.7527 |

Additionally, we found that manually tracing these images took an average of approximately 15 minutes per image, whereas, once trained, the algorithm reduced this time to about 1 minute per image. This effectively speeds up our data segmentation process by a factor of 15. Finally, we note that in this work, we chose not to compare this approach to U-Net [24], a popular CNN model for segmentation, because previous studies have shown that CNN-based region growing outperforms U-Net in terms of retinal vessel segmentation [11] and leaf vein segmentation [10], both of which are similar to dendrite tracing in this study.

### 4.2 Automated dendrite categorization

To evaluate the reliability of the automated dendrite categorization, we compared the algorithm’s predictions with manual corrections across a dataset of 41 segmented PVD images. Performance was quantified using the Jaccard index (intersection-over-union), computed both on a per-image basis and across the pooled dataset (Table 2). Both the per-image and global Jaccard indices were consistently high, exceeding 0.80 for all dendritic categories. This indicates strong agreement between automated and manual classifications across the dataset. Notably, performance was particularly high for fourth-degree dendrites, reflecting the robustness of the algorithm in identifying these distal branches. Together, these results demonstrate that the automated pipeline achieves reliable classification across dendritic classes and generalizes well across different samples. As *C. elegans* age or are subjected to environmental stressors such as hypoxia, cold shock, or osmotic stress, the stereotypical menorah-like architecture of the PVD neuron undergoes structural alterations. To assess whether our algorithm could maintain robust performance under these conditions, we assembled a dataset of images spanning multiple developmental stages (with day 1 defined as the first day following the L4 molt) and compared algorithmic outputs against manually curated ground-truth annotations.

**Table 2.** Jaccard indices for automated dendrite categorization compared to manual annotations. Values are shown as the mean across images and as global indices aggregated across the validation dataset (41 images).

| Category | Per-image mean | Global dataset |
| --- | --- | --- |
| First degree | 0.833 | 0.829 |
| Second degree | 0.849 | 0.845 |
| Third degree | 0.835 | 0.830 |
| Fourth degree | 0.932 | 0.934 |
| Axon | 0.832 | 0.898 |

We observed that with advancing age, the primary (first order) dendrite often developed pronounced bends that produced zigzag patterns. These distortions displaced the dendrite from its central axis and brought it closer to third-order branches, thereby increasing the uncertainty of classification between these categories and divergence from manual annotation (Figure 6A). Such bending also affected the alignment of second- and fourth-order dendrites, which in older worms frequently deviated from their typical orthogonal arrangement and instead formed sharp, irregular angles. These effects were most prominent in seven-day-old animals. In addition to these angular distortions, aged neurons frequently exhibited ectopic branching that did not conform to the canonical PVD morphology, increasing noise for the grouping of axial segments, which reflects a wider trend of age-associated neurite sprouting in *C. elegans* [7,8,25]. For example, we observed branches arising from the primary dendrite that ran parallel to it as opposed to the typical perpendicular second-order branches, as well as tertiary dendrite branches that emerged in atypical positions between the canonical third-order branches and the primary dendrite (Figure 6B). To quantify algorithmic robustness under these conditions, we evaluated performance using the Jaccard index in the developmental stages. As anticipated, precision declined with age in all dendrite categories, with the steepest reductions observed for second-order dendrites and axons (Figure 6C). Notably, even manual annotation becomes increasingly challenging under these conditions, suggesting that part of the observed disagreement reflects intrinsic ambiguity between morphological categories rather than purely algorithmic error.

**Figure 6.**
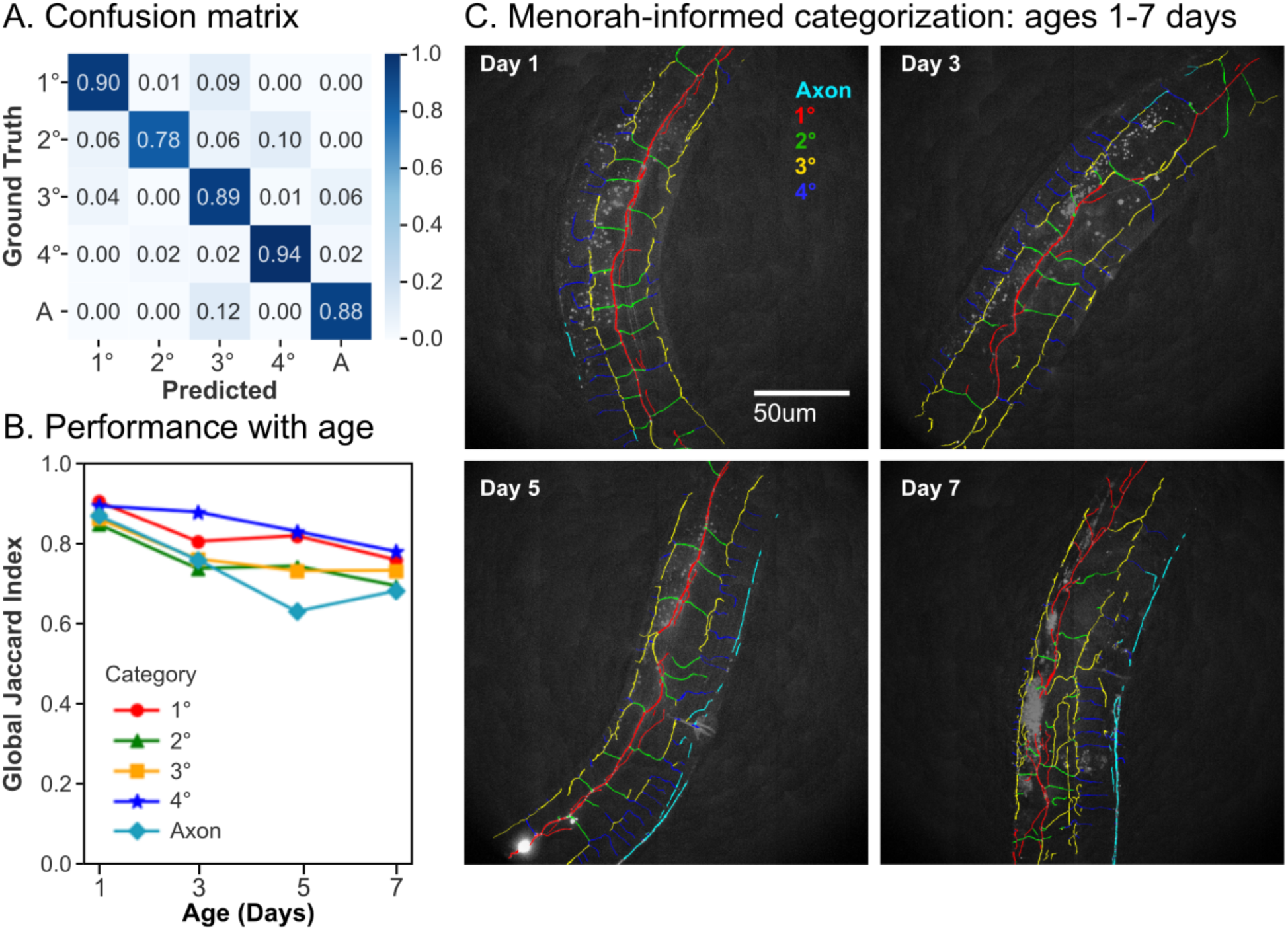
Evaluation of menorah-informed dendrite categorization with worm age. (A) Confusion matrix showing classification accuracy for first- through fourth-order dendrites [1–4°] and axons [A]. High diagonal values indicate robust performance across categories, with most misclassifications occurring between parallel dendrite orders. (B) Global Jaccard Index across developmental time points (Day 1, 3, 5, and 7). While overall accuracy remains high, performance declines with age, particularly for second-order dendrites and axons. (C) Example images of worms at different days of adulthood.

#### 4.2.1. Differences in beading patterns arise from branch-order-specific susceptibility

In agreement with previous observations that distinct beading patterns emerge under different stressors (Figure 7), we confirmed that the morphological patterns of dendritic beading differ markedly between neurons from aged worms and those subjected to acute cold shock [8]. Saberi-Bosari et al. demonstrated that while both aging and cold shock increase the number of bead-like protrusions along PVD dendrites, the spatial distribution of those beads diverges [8]. One of the most salient quantitative distinctions between aging-induced and cold shock-induced beading was the opposing trends in average inter-bead distance. In aged neurons, inter-bead spacing decreased over time, consistent with a progressive accumulation of beads that cluster along dendrites. In contrast, cold shock produced the opposite effect: as the severity of the cold stress increased, beads became more widely spaced, resulting in elevated average inter-bead distances relative to controls. The divergent trends in bead spacing raised mechanistic questions about the underlying dendritic architecture that might drive these phenotypes. To explore hypotheses surrounding structural modifications, we applied our automated dendrite tracing algorithm to characterize the topology and branch structure of the PVD arbor in aged versus cold-shocked animals. This analysis aimed to determine whether changes in overall dendritic length or pruning of distal segments could account for the observed shifts in bead distribution.

**Figure 7.**
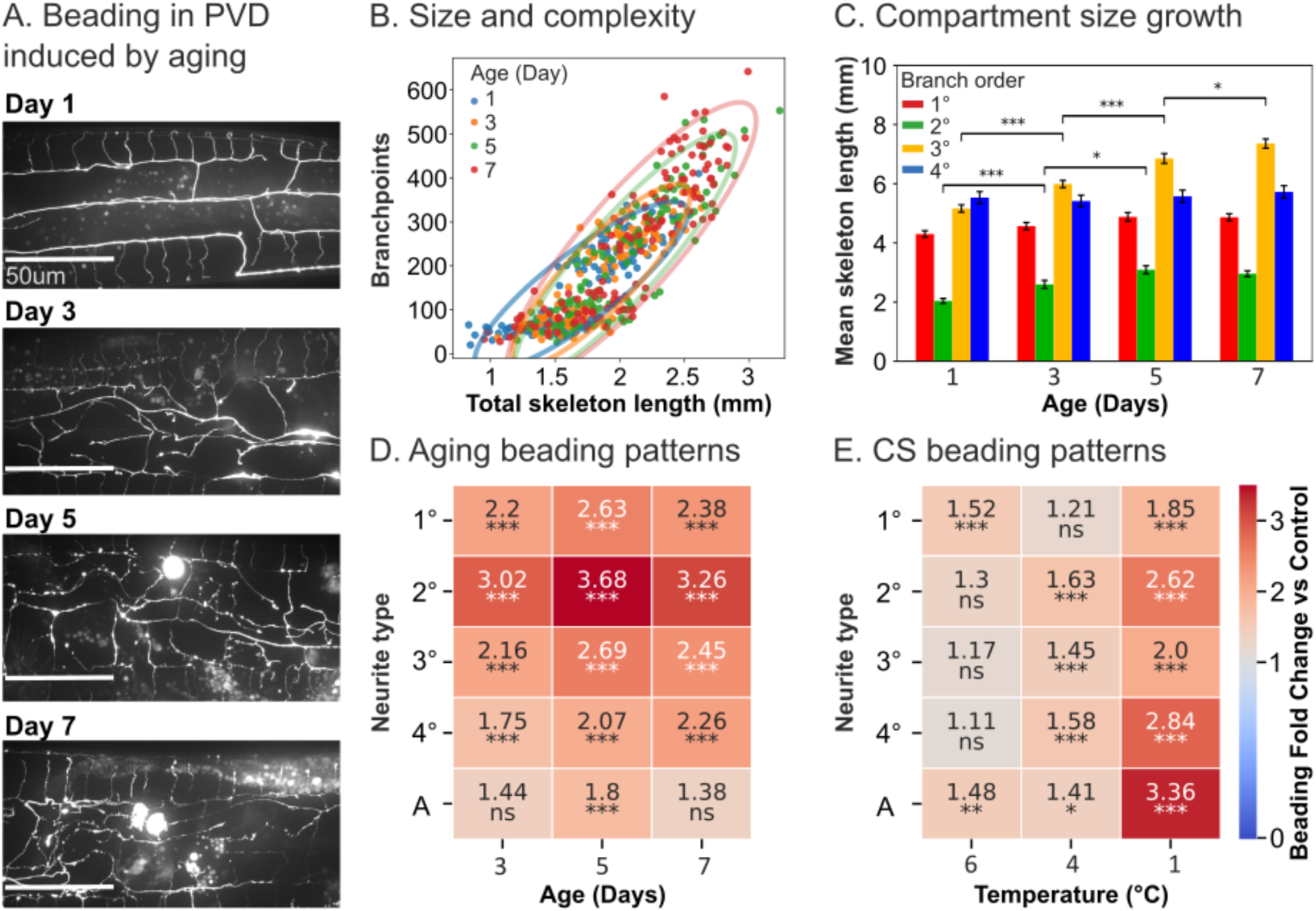
Age-dependent morphological changes of the PVD dendritic arbor and neurite-specific beading behavior. (A) Example images of sections of PVD increasingly diverting structurally and showing beading. (B) Relationship between the number of PVD dendritic branch points per worm and total dendritic skeleton length across age groups. (C) Changes in mean dendritic segment length with aging, stratified by position in the menorah structure, statistical comparisons were performed exclusively between branches of the same order across contiguous dates. (D) Temporal progression of beading abundance across different PVD neurite segments: first- through fourth-order dendrites [1–4°] and axons [A]. (E) Beading propensity of PVD neurites following cold shock treatment. Statistical significance was assessed by pairwise Welch’s two-sample t-test. Significance levels: \**p* < 0.05, \*\**p* < 0.01, and \*\*\**p* < 0.001; P-values were Bonferroni-corrected for multiple comparisons.

First, we quantified branch point abundance to assess whether decreases in average inter-bead distance during aging were driven by pruning of more distal dendrites. Under a pruning hypothesis, one would expect a reduction in the number of branch points as distal segments are lost. However, our measurements revealed the opposite pattern: older worms exhibited a net increase in both total skeleton length and branching points (Figure 7A-B). This finding indicates that distal pruning is unlikely to be the primary driver of reduced inter-bead spacing with age, suggesting instead that the dendritic arbor remains extended and even elaborated in aged neurons.

Because branch point counts alone were insufficient to determine whether aging preferentially affects specific segments of the dendritic tree, we employed a dendrite categorization algorithm to examine average segment lengths across orders of branching. In line with the increased branching observed, segments of all orders either trended toward increased length or remained unchanged with age (Figure 7C). Notably, third-degree branches showed a consistent increase in length, potentially reflecting reduced self-avoidance between neighboring branches of similar order (Figure S1A). This pattern suggests that age-related changes in dendrite geometry do not entail simple truncation of higher-order branches but rather a complex rearrangement of segment lengths.

Given that alterations in dendritic abundance and geometry did not fully explain the age-dependent decrease in inter-bead distance, we next assessed bead distribution relative to branch order. We calculated fold-changes in bead density (number of beads per unit length) for each neurite type, against measurements from day-1 adults (Figure 7D). In aged animals, second order branches displayed relatively high increases in bead density compared to the other compartments, indicating a greater susceptibility to age-related beading (Figure 7D). In contrast, cold shock induced a widespread increase in bead density across all branch types, with second- and fourth-degree branches becoming increasingly vulnerable with cold stress intensity (Figure 7E). This pattern under cold stress aligns with prior work showing that the spatial dispersion of beads is stress intensity-dependent and highlights mechanistic differences between aging and acute stress responses in PVD neuronal morphology [8].

As described earlier, the traditional menorah-based classification fails to accurately capture the identity of dendritic segments that deviate from canonical PVD organization during aging. Consequently, we evaluated whether our morphometric findings remain robust when implementing a branching-informed (parent-child) hierarchical classification. We first tested whether the distribution of average segment lengths mirrored the patterns observed with the traditional menorah-aware method (Figure 8A). First- and second-degree branches exhibited a distinct distribution between the two methods. Specifically, second-degree branches encompassed a greater total length than first-degree branches under the branching-informed framework, which was not the case in the menorah-aware method. This divergence is likely driven by the algorithm’s ability to correctly identify axially oriented second-degree branches that do not give rise to a canonical menorah unit. Additionally, a substantial portion of the total dendrite length was classified as fifth-degree or higher starting from Day 1 of adulthood, with these higher-order branches contributing increasingly to the total arbor length as the animals aged.

**Figure 8.**
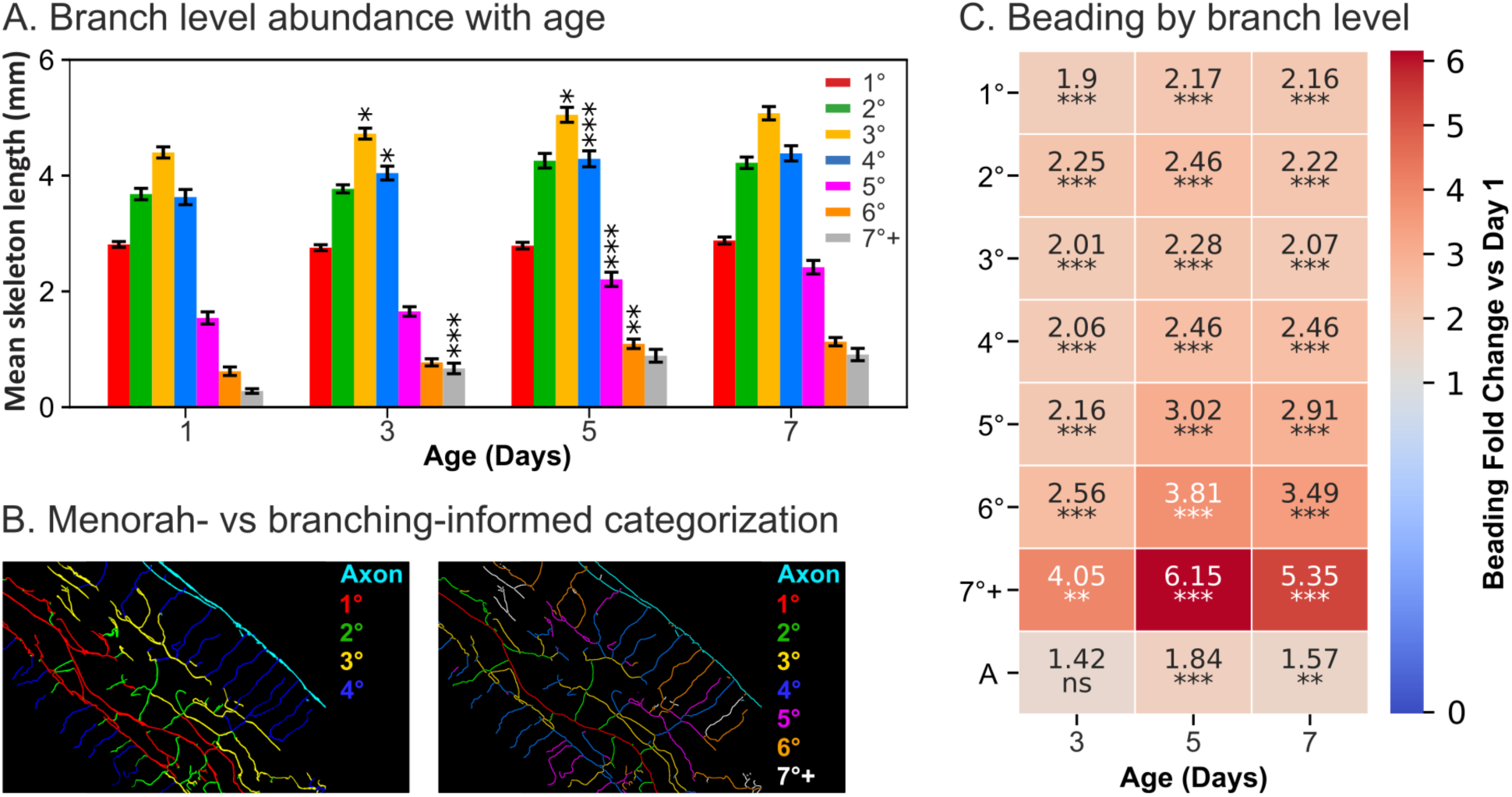
PVD aged phenotype is better captured by unconstrained branching degree analysis. (A) Mean length per worm per branch order, statistical comparisons were performed exclusively between branches of the same order across contiguous dates, asterisks indicate significance of the change with respect to prior age. (B) Example of differences in categorization for aged worms between models. (C) Temporal progression of beading propensity across different PVD neurite types during aging categorized by branching events. Statistical significance was assessed by pairwise Welch’s two-sample t-test. Significance levels: \**p* < 0.05, \*\**p* < 0.01, and \*\*\**p* < 0.001; P-values were Bonferroni-corrected for multiple comparisons.

When analyzing age-dependent beading propensity within this branching-informed framework, second-degree branches did not exhibit a noticeably high vulnerability to beading. Instead, elevated beading propensity was concentrated within segments associated with extended chains of successive branching events. This finding stands in sharp contrast to the traditional menorah-aware paradigm, which identified second-degree segments as the most vulnerable. This discrepancy is mostly explained by the misclassification of degenerate segments occupying the interstitial space between first- and third-degree branches (particularly in aged worms). In the menorah-informed framework, these segments are mostly designated as second-degree by default, regardless of age-induced structural distortion. Conversely, the branching-informed algorithm reveals that many of these interstitial segments are highly branched, higher-order structures (Figure 8B). Together these observations provide an explanation to the reduced inter-bead distances with age, as the most vulnerable segments tend to be highly branch segments in the interstitial space. They also underscore a critical limitation of categorizing processes by relative spatial position rather than by their sequential lineage of growth.

## 5. Discussion

In this work, we developed and validated an automated phenotyping pipeline that combines CNN-based region growing with topology-based dendrite categorization to enable high-resolution morphological analysis of the *C. elegans* PVD neuron. This approach addresses key bottlenecks in neuronal image analysis, particularly for highly branched and structurally complex dendritic arbors that defy accurate segmentation by conventional thresholding tools. By preserving the spatial contiguity of traced neurites, our CNN-guided region growing framework minimizes manual intervention while maintaining high segmentation accuracy across heterogeneous imaging conditions. Importantly, this pipeline enabled the precise quantification of degeneration phenotypes by branch order across two distinct categorization paradigms, offering novel mechanistic insights into how sub-compartments of the dendritic arbor selectively respond to stressors that affect neuronal morphology.

The automated categorization of dendritic segments based on their positions within the canonical menorah structure permitted a systematic investigation of morphological changes during aging and stress. High alignment with manually curated annotations demonstrates that our topology-based classification framework generalizes effectively across individuals and diverse orientations when the baseline menorah architecture is well preserved. Nevertheless, we observed a gradual decline in classification performance with advancing age, particularly among second-degree dendrites and axons. Rather than reflecting a systemic algorithmic limitation, these misclassifications capture genuine biological disorganization. Aging PVD neurons frequently exhibited primary dendrite bending, ectopic branching, and striking deviations from the stereotyped orthogonal branching pattern, consistent with previous reports [6–8]. Such structural breakdown disrupts the stereotypical spatial relationships required for geometric degree assignments, likely mirroring a progressive loss of self-avoidance mechanisms and cytoskeletal regulation. Thus, declining algorithmic precision itself serves as an indicator of morphological deterioration. To circumvent this structural dependency, we developed an alternative categorization strategy that operates independently of spatial coordinates, relying instead on identifying the primary branch and sequentially classifying subsequent segments based on their localized parental continuation or branching transitions.

A central biological contribution of this study lies in elucidating the discrete mechanisms underlying the distinct dendritic beading patterns induced by chronic aging versus acute cold shock. Consistent with prior analyses, we confirmed that aging results in progressively compressed inter-bead distances, whereas cold shock does not (Figure S3). These opposing macroscopic trends raised the possibility that large-scale structural remodeling, such as the pruning of distal dendrites, might account for the altered bead spacing in aged animals. However, quantitative tracing revealed that aging was paradoxically accompanied by an increase in total dendritic length and a higher abundance of branch points, effectively ruling out pruning as a dominant mechanism. Instead, the PVD arbor remains extended and unexpectedly elaborate late into the lifespan.

Further analysis of segment lengths across individual branching orders demonstrated that dendrites of all degrees either elongated or remained stable with age, with third-degree branches exhibiting the most consistent growth. This expansion may reflect a progressive decay of self-avoidance regulatory cues between parallel processes, allowing adjacent branches of similar orders to encroach upon one another’s territory. While these spatial shifts contribute significantly to the crowded architecture of the senescent arbor, they do not directly account for localized changes in bead spacing. Notably, initial bead density analysis using the traditional menorah framework suggested that lower-degree branches, particularly second-order dendrites, accumulated beads at a disproportionately high rate. However, our alternative branching-informed framework revealed that the apparent vulnerability of second-degree processes was an artifact of position-based classification. Age-dependent beading susceptibility is not specific to second-degree segments, but is instead fundamentally dictated by a segment’s position along successive branching events. Crucially, mapping these intricate, distribution patterns is unfeasible through visual inspection; hand-annotation of thousands of crowded segments introduces severe investigator bias and prohibitive time constraints, necessitating an automated analytical approach.

In contrast to the compartment-specific vulnerability seen in aging, acute cold shock produced a widespread, distinct degenerative pattern. Bead density increased uniformly across all branch orders, with severity-dependent shifts determining which dendritic populations were most heavily impacted. While moderate cold stress primarily localized to second-degree branches, extreme thermal stress extended this structural vulnerability outward to fourth-degree distal dendrites. This progressive, synchronized involvement of the entire dendritic arbor likely accounts for the increased global dispersion of beads observed under acute stress conditions. Taken together, these phenotypic signatures indicate that chronic aging and acute cold shock engage fundamentally distinct neurodegenerative pathways: aging preferentially compromises specific dendritic sub-compartments defined by extensive branch-generation history, whereas cold shock triggers a systemic, global structural failure that escalates symmetrically with stress intensity.

The preferential beading of highly branched dendritic segments during aging likely reflects localized deficits in structural or metabolic stability. Distal branches are significantly thinner than primary dendrites yet sustain substantial cytoskeletal loads and topological complexity, potentially rendering them exceptionally sensitive to disruptions in microtubule integrity, actin dynamics, or local plasma membrane tension [26]. Prior studies have linked axonal and dendritic beading to cytoskeletal destabilization and mechanical stress, supporting a model where the chronic, age-dependent weakening of structural support manifests earliest within these hyper-branched compartments [27]. Additionally, the observed growth and crowding of third-degree branches may further exacerbate local stress, promoting bead formation through altered tension or impaired transport. In contrast, the dose-dependent expansion of beading across dendrite orders under cold shock is consistent with acute perturbations such as rapid cytoskeletal depolymerization, membrane rigidity changes, or ionic imbalance [1,28,29]. Previous studies have identified that changes in temperature affect microtubule stability triggering rapid depolymerization at 4°C [30]. The shift from medium-order vulnerability at moderate stress to distal branch involvement at higher stress suggests a cascading failure model, where initial damage propagates outward as cellular homeostasis collapses.

Future work could integrate longitudinal imaging of individual neurons to track the temporal emergence of beading and morphological rearrangements, potentially enabling predictive modeling of degeneration onset. Coupling this framework with fluorescent reporters of cytoskeletal integrity, calcium signaling, or metabolic stress would further clarify the mechanisms driving compartment-specific vulnerability. Moreover, the generalizability of CNN-based region growing suggests potential applications to other neuronal types and biological network structures.

## 2.7 Acknowledgements

John Lagergren’s current affiliation is Oak Ridge National Lab, but this work was performed during his studies at North Carolina State University.

